# A bioinformatic panel to interrogate thousands of ExAC variants with minor reference allele that are missed by conventional variant calling

**DOI:** 10.1101/093450

**Authors:** Mahmoud Koko, Mohammed O. E. Abdallah, Mutaz Amin, Muntaser Ibrahim

**Affiliations:** Department of Molecular biology, Institute of Endemic Diseases, University of Khartoum, Khartoum,Sudan.; Department of Neurology and Epileptology, Hertie Institute for Clinical Brain Research, Tuebingen,Germany.; Department of Biochemistry, Faculty of Medicine, University of Khartoum, Khartoum, Sudan.

**Keywords:** minor reference alleles, human genome, ExAC, variant calling

## Abstract

In variation sites with minor reference alleles, overlooking the detection of homozygous reference genotypes results in inadequate identification of potential disease variants. Current variant calling practices miss these clinically relevant alleles warranting new approaches. More than 26,000 Eome Aggregation Consortium (ExAC) variants have a minor reference allele including 44 variants with known ClinVar disease alleles. We demonstrated how the current variant calling standards miss homozygous reference disease variants in these sites. We developed a bioinformatic panel that can be used to screen these variants using commonly available variant callers. We provide here a simple strategy to screen potential disease-causing variants when present in homozygous reference state.

## Introduction

The basic units in many if not most genomic studies are “genetic variations”. Although the name itself implies variation between individuals, the choice of a single reference is a practicality imposed by the need for a common ground to base the genomic analysis. The alignment of sequences to this reference is tolerant to variations (Li H, 2013). The basic concept of variant calling is, on the contrary, contrasting the sequence reads against a reference to detect these mismatches (McKenna et al., 2010). Choosing the best possible combination of reference alleles to each and all variation sites in the human genome is thus critical to the frictionless calling, annotation and prioritization of variations (Dewey et al., 2011; Ibrahim et al., 2016). A debate is still in place whether the major alleles or ancestral alleles constitute this alleged best. The fact remains, however, that many variation sites in the human genome harbor an allele denoted as a reference – with a frequency lesser than other alternate allele(s) (Magi et al., 2015).

These sites, called hereafter “minor reference alleles”, are challenging during variant calling. Reference alleles with low allele frequencies are sites of mismatch for the majority of human population, appearing frequently as homozygous or heterozyous variants with high alternate allele frequency that is more than 0.5. These sites are usually filtered upon prioritization in disease studies. A lesser fraction of the population will have a homozygous rare allele that will pass undetected because both alleles match the human reference and thus “do not have a variant”. Keeping in mind that most studies look for alleles with lesser frequencies in the human genome, these sites bearing homozygous variants – that could relate directly to the phenotype – will be missed during conventional variant calling.

Previous reports have proposed some strategies to approach such reference alleles. Dewey et al (2011) used a modified version of the human genome as a “major allele” population specific reference. Magi et al. (2015) studied extensively the count and annotation of “rare” reference alleles in 1000 Genomes data and developed a specialized variant caller, RAREVATOR, based on GATK UnifiedGenotyper. We studied the latest release of ExAC data as a more extensive source for coding variants (Lek at al., 2016) to find out the extent of minor reference alleles in the human genome in different ExAC populations and their implication on variant calling.

We propose here a simple flexible strategy to interrogate these variations based on a bioinformatic panel of minor reference allele sites. This approach can be implemented in principle by targeted calling in specified regions which can be specified based on global or population-based ExAC allele frequencies, allowing the detection of homozygous reference alleles. We discuss other aspects affecting variant calling in minor reference allele sites.

## Minor reference alleles in ExAC

We defined minor reference allele variants as variants with non-reference allele(s) frequency of 0.5 (reference alleles with a frequency lesser than the total frequency of the other observed alleles in the same variation site). Although “minor allele” refers in general to the second-most but not necessarily the least frequent allele, and does not require a frequency threshold, this practical definition used for the purpose of this report allowed the identification of reference alleles present in less than half of ExAC sample cohort without a bias in multi-allelic sites. We based this arbitrary threshold on the observation that ClinVar alleles are seen with frequencies as high as 0.46 (rs3733402, ClinVar variation ID 12037). We found that the latest ExAC release 0.3.1 VCF contained 26537 variants with a reference allele frequency less than 0.5 (minor reference allele). In total, 26529 variants were annotated with rsIDs from the latest dbSNP release 147, including three variants with an old rsID that was merged in a new record (with possibly new coordinates). Eight variants were not found in dbSNP. Around 1% of these variants (2763 variants) were rare variants (AF < 0.01). The the characteristics of these variants are shown in table (1). Out of this pool, a group of variants had a ClinVar significance of (likely)pathogenic, risk factor, association, or drug response allele (table 2).

**Table (1):** summary of minor reference alleles in ExAC database: Multi-allelic:

| Variants, sites | bi-allelic | multi-allelic | total |
| --- | --- | --- | --- |
| SNVs | 20065 | 4448 | 24513 |
| Indels | 890 | 2286 | 3176 |
| SNVs & Indels | NA | 1152 | 1152 |
| Total | 20955 | 5582 | 26537 |

**Table (2):** ClinVar variants with minor reference alleles. This subset contains only variants with these annotations: pathogenic, likely pathogenic, risk factor, drug response, or association. Allele frequencies are reported from ExAC. The variants are given in order of minor allele frequencies.

| rsID | REF |  | Minor Allele |  |
| --- | --- | --- | --- | --- |
|  | hg19 | hg38 | Minor | MAF |
| rs1169305 | A | A | A | 0.004 |
| rs4784677 | C | C | C | 0.006 |
| rs6025 | T | C | T | 0.02 |
| rs283413 | A | C | A | 0.02 |
| rs4918 | G | G | G | 0.03 |
| rs820878 | T | T | T | 0.03 |
| rs2476601 | A | A | A | 0.07 |
| rs450046 | C | C | C | 0.08 |
| rs12021720 | T | T | T | 0.09 |
| rs6003 | C | C | C | 0.13 |
| rs1154510 | T | T | T | 0.15 |
| rs1676486 | A | A | A | 0.2 |
| rs1051931 | A | A | A | 0.2 |
| rs7076156 | A | A | A | 0.21 |
| rs1366813 | T | C | C | 0.22 |
| rs1801265 | G | A | G | 0.23 |
| rs1799983 | T | T | T | 0.25 |
| rs497116 | T | T | T | 0.25 |
| rs2227564 | T | T | T | 0.25 |
| rs1800858 | A | A | A | 0.26 |
| rs20541 | A | A | A | 0.27 |
| rs2278619 | A | A | A | 0.27 |
| rs1042714 | G | G | G | 0.3 |
| rs4917 | T | T | T | 0.32 |
| rs10774671 | G | G | G | 0.33 |
| rs1061170 | C | C | C | 0.33 |
| rs6280 | C | C | C | 0.34 |
| rs10010131 | A | A | A | 0.34 |
| rs1494555 | G | G | G | 0.35 |
| rs5219 | T | T | T | 0.35 |
| rs1260326 | T | T | T | 0.36 |
| rs16891982 | C | C | C | 0.36 |
| rs1494558 | T | T | T | 0.37 |
| rs1341667 | T | T | T | 0.38 |
| rs5882 | G | G | G | 0.4 |
| rs853326 | A | A | A | 0.42 |
| rs2073711 | A | A | A | 0.43 |
| rs6166 | C | C | C | 0.43 |
| rs1131454 | G | G | G | 0.44 |
| rs237025 | G | G | G | 0.44 |
| rs2303067 | A | A | A | 0.45 |
| rs699 | A | A | A | 0.45 |
| rs1815739 | T | C | T | 0.45 |
| rs3733402 | G | G | G | 0.46 |

### Updated references in assembly 38

Lift-over to the new assembly hg38 indicated that 1212 known variants had an updated reference allele. The remaining sites (after removal of updated references and remapping to nonconventional chromosomes) were used to construct an updated panel with hg38 coordinates (additional file 1).

### Minor Reference Alleles Panel

We used all these ExAC minor reference variants to create a bioinformatic panel with hg19 coordinates to force variant calling of homozygous reference variants. We constructed panels for all minor reference allele sites and rare sites (additional file 1). Using either of these panels, the combination of targeted allele genotyping, and filtering for homozygous reference alleles allows to screen homozygous reference variants.

### Population-based minor reference alleles

The allele frequencies for many variants tend to show a spectrum of variation across populations, especially when selection, drift, or founder effects are notable. We show in table (3) the number of variants that had a minor reference allele in each population of the ExAC populations. When examining samples from a specific population, the use of panels based on these estimates might be more appropriate (additional file 2).

### Targeted Variant Calling of MRA sites

Exome data sets for Genome In A Bottle sample HG001 (NA12878) were used for the purpose of testing this panel strategy. We used the minor reference allele panel VCF as “alleles” input to GATK HaplotypeCaller to limit the genotyping to the panel alleles. Slightly more than 1000 homozygous reference variant calls in minor reference allele sites were seen at a depth threshold of 10x (additional file 3). Around 800 variants were seen at a depth threshold of 30x with high genotype qualities. As site qualities (QUAL field in VCF) are set to zero at reference sites, we evaluated the RGQ (reference genotype qualities) using the same panel as “region” input for GATK HaplotypeCaller. We found that the majority of panel sites showed good depth and very high RGQ (additional file 3).

## Discussion

One major challenge that faces the genomic revolution is how to perform an accurate analysis to provide high confidence results. In genome and exome sequencing experiments, variant callers are crucial to this challenge. Many techniques and recommendations were developed to improve the calling accuracy. For instance, “GATK Best Practices” is one contemporary way of giving a reliable and validated road-map for variant calling (DePristo et al., 2011). The essential concern has always been to reduce the number of false calls, whether false positives or negatives. It is a common knowledge that exome and genome sequencing will miss some variants – however small their number might be – that are poorly enriched or sequenced and thus do not show enough information or quality to enable adequate calling. Nonetheless, with acceptable quality and coverage, one assumes that variants would be reliably identified. It might seem a surprise that a considerable number of variants are discarded as routine despite being very reliable and valuable. The notion that a sample “has this variant” or “does not have that variant” is in itself misleading. In a sense, every human genome has all variants but a different combination of alleles. Limiting variant calling to differences is thus an underestimation; homozygous reference variants are usually treated the same way as the remaining parts of the genome where no variations are observed. The recent versions of GATK started to report reference genotype qualities to help highlighting these sites.

Magi et al. (2015) have recently addressed the presence of “rare” reference alleles in the 1000 Genomes project data that are missed by standard variant calling practices, proposing the name “rare reference alleles”. To our knowledge, RAREVATOR is the only variant caller specifically designed to detect rare reference alleles (Magi et al. 2015). It is based on the widely used GATK UnifiedGenotyper. However, HaplotypeCaller is a more reliable haplotype based caller for relatively small sample numbers (Van der Auwera et al., 2013).

### Minor reference allele targeted calling

A subset of ExAC variants with minor reference alleles can be used as described in this report in parallel to the usual variant calling practice to force calls only on such positions followed by filtering for homozygous reference variations. This provides a simple and flexible method to screen possible relevant variants. For instance, this panel can be intersected with BED intervals of enriched targets to get calls on sequenced regions only (see methods). It can be limited in the same manner to a list of genes, coding regions or loci under study. In principle, this can be used with all variant callers that support defining variant calling regions. As the number of variants is limited, it does not require any special computational setting more that what is used to do the original calling and can be run on previously aligned samples with the same variant caller as shown in HG001. More importantly, the panel can be customized in a population-oriented way to reflect reference allele frequencies in a specific population by using population-based ExAC frequencies for panel construction. It could be constructed from any VCF with AF data including local cohorts. Although one can opt to report all variant sites using dbSNP VCF, the minor reference panel approach will provide clear advantages. Besides being considerably lesser in number and thus easier to call, it insures that only minor reference alleles are included, and accordingly, all homozygous reference alleles are of interest. With the inclusion of more samples in ExAC project, many 1000 Genomes minor alleles has been shown to have a higher frequency as previously reported. Using population allele frequencies to build a panel, it is possible to get a list of variants where the human genome harbors minor reference alleles for the specific population. When combined with standard calls, the identification of homozygous reference alleles based on ExAC frequencies for the population under study will help to improve the identification of variants in a population based fashion.

Other possible solutions beside dedicated variant callers and the use of a bioinformatic panel as reported here encompass forced calling at all dbSNP sites, joint calling, and population-based references.

### Variant detection in single and multiple samples

Joint calling of multiple samples has the advantage of forcing variant calls in all samples if one sample has a single non-reference allele. In this manner, it takes only one copy of a non-reference allele in one sample to see homozygous rare reference alleles in all the other samples. Taking in consideration that the non-reference allele is actually common, it is quite likely that it will be there in one of the samples if the cohort is sufficiently large and diverse. Odds get even better as the reference allele shows a lower frequency – the chances of having a heterozygous or homozygous alternative sample then go higher. That being said, one should always keep in mind an important issue of population stratification. Small number of samples from the same population with the same phenotype can have the same homozygous allele that is considered otherwise rare in a global setting and thus escaping joint calling. This panel can be used with joint calling to detect homozygous reference variants in all samples.

In this era of personalized approach to treatment, it is not uncommon to perform an analysis of a single exome sample (reference). In this case, the detection of minor reference alleles is of particular importance. (? actionable variants)..

### Different reference alleles at the same site

The use of population-based references for analysis has been previously suggested (Dewey et al. 2011). However, this might create many inconsistencies in reporting variants and technical difficulties to maintain multiple “human genomes”. The current methods for annotation of variants are dependent on the version of the human genome. Consistency in reference alleles is a prerequisite for bioinformatics tools to perform any sort of comparisons between results. The use of targeted calling at sites of interest complements the advantages of having a single genome by specifically calling problematic variation sites. It offers the opportunity to be updated with every human genome build and incorporates findings from any future large scale genome projects to update the definition of minor reference allele sites, including the expected update of ExAC to 120,000 samples. Even with the current use of a single reference genome, a lot of inconsistencies in reporting alleles are already observed, mostly caused by the conflicting definitions of minor alleles and references, and the tendency to report the non-pathogenic allele as reference. The variant rs16891982 in the SLC45A2 gene is one example. The minor, reference and ancestral allele is C with ExAC frequency of C=0.3597. The 1000 Genomes give a different minor allele (G=0.2750). The ExAC allele frequency ranges from 0.0022 in East-Asian population to more than 0.95 in European populations, with a frequency of 0.1546 in African population. The allele G is reported in ClinVar (NM_016180.4:c.1122C>G (p.Phe374Leu)) to be associated with dark skin pigmentation and protective effect against malignant melanoma. A separate entry is available for the reference allele C as benign (NM_016180.4:c.1122C= (p.Phe374=)). The plus sign '=' indicates that no change has happened, in other words, the reference is the reported clinical allele. Interestingly, this variation site is linked to an OMIM entry where a report by Stacey et al. (2009) associates the allele G with increased risk of squamous and basal skin cancer, referring to allele G as reference. Detection of homozygous reference changes can help normalizing the reporting of variants in relation to one reference genome.

### Allele frequencies, sample size and population stratification

The ExAC data we used in this report provides an additional advantage of a larger sample size (60,706 individuals) compared to the 1000 Genome samples studied previously (Magi et al. 2015) and thus gives a better estimation of coding reference allele frequencies and population based estimates at a higher resolution compared to previous sets. The variant rs1131454, which has been reported as a “risk factor” locus for type 1 diabetes mellitus, is a good example. This variant has a 1000 Genomes allele frequency of A=0.4738 from an allele count of 2,373. On the other hand, the allele frequency in ExAC is A=0.5628, based on an allele count of 68,216, giving a more reliable estimate. Interestingly, it shows a wide population-based variation in alternate allele frequencies ranging from A=0.7033 in Latino population (AC=8131, AN=11562, homozygotes=2919) to A=0.2286 in African population (AC=2376, AN=10394, homozyotes=298). A study cohort from the Latino population would have approximately 10% individuals with G/G genotype. A cohort from an African population would have 60% individuals with G/G reference genotype potentially skipping identification (the actual percentage falls considerably with joint calling as we discuss below). The inconsistency in calling such a variant would seriously affect efforts to validate the effect of this variant.

### Looking for the rare and overlooking the common

Not only rare reference alleles are important in this context. It is not necessarily that an allele has a frequency less than 0.1 to be potentially significant and problematic during calling. Under a simple hypothesis of a detrimental allele effect, some genome variation sites would have a minor “pathogenic” allele: the larger the effect size, the “rarer” the allele gets. Filtration based on small thresholds of allele frequencies is thus one of the established (but not necessarily great) prioritization approaches that will not give the best outcome in non-mendelian or non-penetrant phenotypes. Variants with common alleles are already associated with many clinical conditions. ClinVar database contains many documented variants where the disease related alleles have a frequency well above 0.1 up to 0.4 (table 2). Such variants that usually have a moderate to low effect size are seen in a wider number of individuals in homozygous state compared to rare alleles, thus potentially missed in larger numbers of individuals. Missing such variants will compromise the results of genomic analysis especially when the allele is particularly common in a specific population under study. These sites are also of interest when the disease phenotype does not readily affect the fitness of the individual, or otherwise the effect of multiple alleles exerts the phenotype in a complex inheritance model. Part of the missing heritability in genomic studies is largely attributed to the joint effect of such variants that are “not-rare-enough” to be confidently included but “not-common-enough” to be excluded as a pathogenic allele. In our analysis, we found it more appropriate to pursue the identification of any reference allele that is less frequent than the alternate allele(s). The name “minor reference allele” is thus more descriptive in this regard despite the possible confusion with “minor allele” defined conventionally as the second most common allele in a given population.

## Conclusions

Reporting variants in minor reference allele sites is one example of the hurdles that face the communication of results in genomics. Target calling using a panel of minor reference allele sites will help to scan homozygous minor reference alleles with remarkable flexibility and population based customization.

## Methods

### Minor Reference Allele sites in ExAC

We downloaded ExAC release 0.3.1 VCF file (Lek et al., 2016) and performed a pair-wise comparison of frequencies between reference alleles and all observed alternate alleles as follows: variants with multiple alleles were split using vt tool (Tan et al., 2015) automatically sub-setting allele specific frequencies; alleles and their frequency data from the resulting VCF were filtered based on reference allele frequency cut-off = 0.5 using a custom script written in R (R Development Core Team, 2008) giving all minor reference allele sites in ExAC data in human genome assembly hg19; the results were used to construct a bioinformatic VCF panel for targeted calling; to construct population based panels, the custom script was modified to calculate population allele frequencies from allele numbers and counts. **Testing against GIAB Whole Exome data sets**: We used bam files for sample HG001 (NA12878) publicly available from GAIB to perform targeted allele calling (Zook et al., 2016). The two exomes were sequenced from one sample enriched using *nextrarapid* library on two lanes on Illumina HiSeq2500 platform (Illumina, San Diego, CA, US), aligned using novoalign (Novocraft Technologies, Malaysia), and preprocessed according to GATK best practices recommendations (Base Quality Score Recalibration and Indel Realignment)(McKenna et al., 2010; DePristo et al., 2011; Van der Auwera et al., 2013). We used a subset of the bioinformatic panel that overlapped with the enrichment regions as input to GATK HaplotypeCaller tool in “genotype given alleles” mode to limit genotyping to the panel alleles and in “discovery” mode to obtain reference genotype qualities. **Reference allele discordance between assemblies:** We lifted-over the genomic coordinates of the panel (VCF) using NCBI remapping service to obtain the human genome assembly 38 equivalent coordinates, followed by removing sites with reference update and taking only first pass remapping.

## Additional files

additional file (1) is a zip file containing minor and rare reference allele sites in the latest ExAC release for hg19 and hg38 constructed as described in results and methods. Additional file (2) is a zip file containing population-based panels. Additional file (3) is a zip file containing results of genotype calling and reference genotype qualities in GIAB sample HG001 (NA12878, NIST7035).

## Availability of data and materials

The datasets analyzed in this study are available in the ExAC repository [ftp://ftp.broadinstitute.org/pub/ExAC_release/release0.3.1/] and Genome In A Bottle repository [ftp://ftp-trace.ncbi.nlm.nih.gov/giab/ftp/data/]. The NCBI genome remapping service is available at [www.ncbi.nlm.nih.gov/genome/tools/remap]. Detailed methods, key data sets generated following the analysis, and links to the input files are included in this article and its additional files.

## Authors’ contribution

MK conceived the research idea. All authors contributed to the design. MK performed the bioinformatic analysis with input from MA, MOEA, and MI. The manuscript was written by MK with intellectual input and revisions from MA, MOEA, and MI. All authors read and approved the final manuscript.

## Authors’ information

Muntaser Ibrahim (PhD.) is a professor of genetics and PI at the Institute of Endemic Diseases, University of Khartoum. Mahmoud E. Koko (MBBS, M. Sc.) is a research fellow at the Institute of Endemic Diseases (IEND), University of Khartoum (U of K) and currently a PhD student at the Experimental Epileptology Research group, Hertie Institute for Clinical Brain Research (Tuebingen, Germany). Mohamed O.E. Abdallah (MBBS, M. Sc.) is a lecturer of immunology and microbiology, International University of Africa, Khartoum, and a PhD student at the Bioinformatics and Genomics research group, IEND, U of K. Mutaz Amin (MBBS, M. Sc.) is a lecturer of biochemistry and genetics, Faculty of Medicine, U of K.

## Competing interests

The authors declare that they have no competing interests.

## Ethics approval and consent for publication

Not applicable.

## Open Access

This pre-print is distributed under the terms of the Creative Commons Attribution 4.0 International License (http://creativecommons.org/licenses/by/4.0/), which permits unrestricted use, distribution, and reproduction in any medium, provided you give appropriate credit to the original author(s) and the source, provide a link to the Creative Commons license, and indicate if changes were made.

